# Distinct satellite DNA composition between core and germline restricted chromosomes in *Bradysia (Sciara) coprophila*

**DOI:** 10.1101/2025.04.15.648937

**Authors:** Anne Kerrebrock, Jullien M Flynn, Robert B Baird, Christina N Hodson, Laura Ross, Yukiko M Yamashita

## Abstract

Programmed DNA elimination (PDE), a phenomenon wherein cells eliminate a subset of genetic material during certain stages of development, is observed in a broad range of organisms. The fungus gnat *Bradysia* (formerly *Sciara*) *coprophila* undergoes a series of PDE events during their development, including elimination of germline-restricted chromosomes (called L chromosomes) in soma and elimination of paternal chromosomes during male meiosis. However, the underlying mechanisms of this phenomenon are poorly understood. Highly repetitive satellite DNA, which often shows chromosome specific distribution, is a possible candidate for sequences involved in PDE. In this study, we utilized recent genomic data and genome assemblies to identify new satellite DNA sequences of *B. coprophila*. Through characterization of satellite DNA distribution on chromosomes, we found that the X and autosomes do not share centromeric satellite DNA sequence with the L chromosomes. We further provide the cytological evidence that confirms a recent finding based on the genome assembly that there are two distinct L chromosomes that were not previously distinguished cytologically. Together, our work lays a foundation for future studies to explore the possible connection between satellite DNA and the mechanism of PDE in *B. coprophila*.

## Introduction

Programmed DNA elimination (PDE) is a phenomenon by which cells eliminate a subset of genetic material during certain stages of development. PDE may occur via chromosome elimination, wherein the entire chromosome(s) are lost, or chromosome diminution, wherein parts of chromosomes are deleted (reviewed in Dedukh and Krasikova, 2022; Drotos et al., 2022; Zagoskin and Wang, 2021). Some examples of PDE occur in the germline, where, for example, chromosomes of a particular parental origin are eliminated (reviewed in Herbette and Ross, 2023). Other examples occur during embryogenesis and result in elimination of genetic material from the soma but retention in the germline, e.g. in hagfish, lamprey, nematode and songbird (Nakai et al., 1991; Smith et al., 2009; Tobler, 1986; Torgasheva et al., 2019). The biological significance of PDE remains largely a mystery: why do cells eliminate DNA of a certain parental origin? What is the function of eliminated DNA in the cells that retain them? It has previously been speculated that germline-limited genetic material may have important germline-specific functions (Bryant et al., 2016; Feng et al., 2017; Kinsella et al., 2019; Marlétaz et al., 2024; Wang et al., 2017) or that PDE is a mechanism to eliminate excess repetitive DNA from somatic cells, including satellite DNA (Kubota et al., 2001; Timoshevskiy et al., 2019; Wang et al., 2017) and transposable elements (Sun et al., 2014) to avoid deleterious effects on cell growth or development. PDE is also involved in sex determination and dosage compensation in certain organisms (Hodson and Ross, 2021; Smith et al., 2021; Streit et al., 2016). Understanding the function or significance of PDE and/or DNA that is eliminated via PDE requires the knowledge of the mechanisms that underpin this phenomenon.

The fungus gnat *Bradysia* (formerly *Sciara*) *coprophila* has long served as a model system to study PDE because they undergo multiple types of chromosome elimination: elimination of germline restricted chromosomes (GRCs, called L chromosomes) from somatic cells during embryogenesis and elimination of paternal chromosomes from male germ cells during meiosis (Figure 1) (reviewed in Gerbi, 2022). Upon fertilization, all embryos in this species start with three X chromosomes, three L chromosomes, and two sets of autosomes (XXX LLL AA), where sperm contribute two Xs, two Ls and one set of autosomes (XX LL A), whereas eggs contribute a haploid genome (X L A) (Figure 1A). *B. coprophila* undergoes four distinct types of PDE: early during embryogenesis, all somatic cells eliminate the L chromosomes (Figure 1A) (Du Bois, 1933). Subsequently, somatic nuclei also eliminate extra X chromosomes: female embryos eliminate one X and male embryos eliminate two Xs, to produce XX females and XO males, respectively (Du Bois, 1933)(Figure 1B). These somatic eliminations of X and L chromosomes occur during a time when the nuclei are actively dividing, and the sister chromatids of eliminated chromosomes lag at anaphase and are unable to segregate to the poles (de Saint Phalle and Sullivan, 1996; Du Bois, 1933). Later in embryogenesis, germ cells eliminate one X and one L chromosome to yield the germline karyotype of XX AA LL in both sexes (Figure1C) (Rieffel and Crouse, 1966). This occurs when the cells are not mitotically dividing (Rieffel and Crouse, 1966), and the chromosomes to be eliminated are extruded from the nucleus in the form of micronuclei (Perondini and Ribeiro, 1997). Finally, the paternally-inherited X and autosomes are eliminated during spermatogenesis as a result of an asymmetric first meiotic division, such that males only transmit chromosomes they inherit from their mothers, along with both copies of the L chromosomes (Figure 1D) (Metz, 1938). Although PDE has been known to occur in *Bradysia* for many decades, the precise cellular and molecular mechanisms remain unknown.

**Figure 1.**
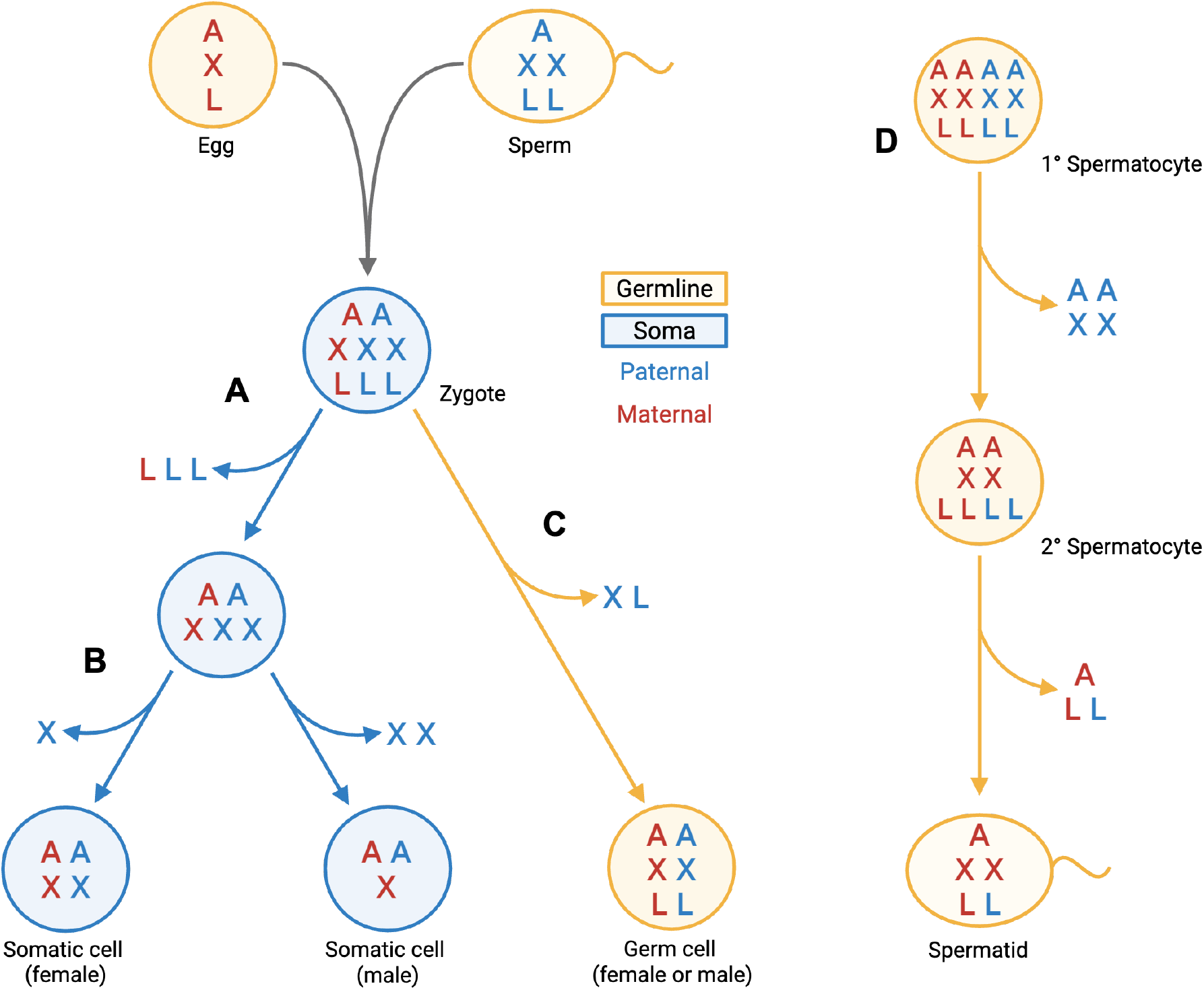
Chromosome eliminations in *Bradysia coprophila* embryos. Zygotes begin with three X chromosomes resulting from fertilization of X eggs by XX sperm. In early embryogenesis, germline-restricted L chromosomes are eliminated from somatic cells (A) shortly followed by X chromosomes (B). The embryo develops as a female or male if one or two X chromosomes are eliminated, respectively. One X and one L are eliminated in the germ cells of both sexes during late embryogenesis (C). In male meiosis, the paternally-inherited X chromosome and autosomes are eliminated in the first meiotic division. In the second meiotic division, the X undergoes nondisjunction, resulting in two X chromosomes in the single spermatid. (D)

Satellite DNA sequences are highly repetitive tandem arrays of non-coding DNA, often constituting a substantial amount of chromosome, forming heterochromatin (reviewed in Garrido-Ramos, 2017; Šatović-Vukšić and Plohl, 2023). Some satellite DNA sequences are specific to individual chromosomes and/or individual species, often serving as useful markers for identifying chromosomes and/or species (Jagannathan et al., 2017; Lohe et al., 1993; Lohe and Roberts, 2000). However, their highly repetitive nature often imposes a challenge in genome assembly, and the location of satellite DNA within the genome is not always clear. In *B. coprophila*, only three families of satellite DNA have been identified thus far (Escribá et al., 2011), and many more satellite DNA sequences are likely yet to be identified. Highly repetitive DNA has been shown to be a target of somatic chromosome elimination in several species including hagfishes, lampreys and parasitic nematodes (Kubota et al., 2001; Timoshevskiy et al., 2019; Wang et al., 2017; Zagoskin and Wang, 2021), thus characterization of satellite DNA in *B. coprophila* may also provide insights into mechanisms of PDE in this species.

In this study, by analyzing recent genome sequencing data of *B. coprophila*, we identified several new satellite DNA families, thus broadening the known repertoire of satellite DNA sequences in this species (Escribá et al., 2011). We found that some satellite DNAs were clustered near the centromeres and ends of chromosomes, while others were broadly interspersed across the genome. Our analysis revealed two interesting features of *B. coprophila* chromosomes. First, we show that a 155 bp satellite DNA, which is found on the centromeres of the core chromosomes (X and autosomes) (Escribá et al., 2011), is absent from the germline-restricted L chromosomes. Second, we found that the two L chromosomes, which were previously not distinguished cytologically, are two distinct chromosomes with distinct satellite DNA compositions. This confirms results from a recent germline genome assembly where L chromosomes were identified due to coverage differences between the contigs of each chromosome (Hodson et al., 2022). Our findings also show that satellite DNA composition is distinct on the germline-limited L chromosomes relative to the core chromosomes, and prompt further study of the potential relationship between satellite DNA and chromosome eliminations in *B. coprophila*.

## Results

### Identification of new satellite DNA families in the *Bradysia coprophila* genome

Previously, three *B. coprophila* satellite DNA repeats were identified by microdissection of the pericentromeric region of the X chromosome, followed by cloning of the microdissected DNA into a plasmid vector (Escribá et al., 2011). This approach identified a 42 bp satellite DNA confined to the proximal heterochromatin on the X chromosome, a 382 bp satellite DNA present on the X, IV and L chromosomes, and a 155 bp satellite DNA (Sccr) at the centromeres of all chromosomes (Escribá et al., 2011). Because these three satellite DNAs likely only represent a small fraction of satellite DNAs of *B. coprophila*, we utilized recent genome sequence data to identify more satellite DNAs.

Using the *B. coprophila* core chromosome (i.e. X and autosome) scaffolds (Urban et al. 2021), we found that tandem repeats make up approximately 5.5% of the genome assembly (a full list of tandem repeats is available in Supplementary file 1). We found 651 satellite DNA repeat sequences which had a unit size between 30-180 bp. These 651 satellite DNA sequences were clustered into 191 satellite DNA families. We selected the eight most abundant families (each family spanning 200 kb to almost 2 Mb in the assembly) to further analyze their distribution within the genome. The total amount of the eight most abundant satellite DNAs summed to 5.6 Mb (Table 1, representative sequences of the repeat units are in Table S1). The eight satellite DNA families were named according to the naming convention recommended by (de Lima, 2022): (Species abbreviation) Sat(abundance ranking)-(unit length), i.e., ‘BcopSat1-145’ indicating the most common satellite family in *B. coprophila* with a 145 bp unit repeat. Among these was the 155 bp ‘Sccr repeat’ family described previously (Escribá et al., 2011), here referred to as BcopSat2-155. The other seven satellite DNA families are newly described, and have unit repeat lengths between 37 bp and 176 bp. Each satellite DNA family contains various minor sequence and length variants, although the repeat divergence landscape shows a high degree of homology (>90% sequence identity) within families (Figure S2). The satellite DNA families BcopSat1-145, BcopSat3-176 and BcopSat4-37 had two abundant sequence variants, which we will refer to as A and B (e.g. BcopSat1-145A and BcopSat1-145B). Two of the previously identified satellite DNAs, with 42 bp and 382 bp repeat units, had low abundance in the assembly (8 and 42 kb total, respectively) (Escribá et al., 2011). Together, these analyses identified highly abundant satellite DNAs in the *B. coprophila* genome, totaling 5.6 Mb (1.7% of the assembly), most of which are identified for the first time.

**Table 1.**
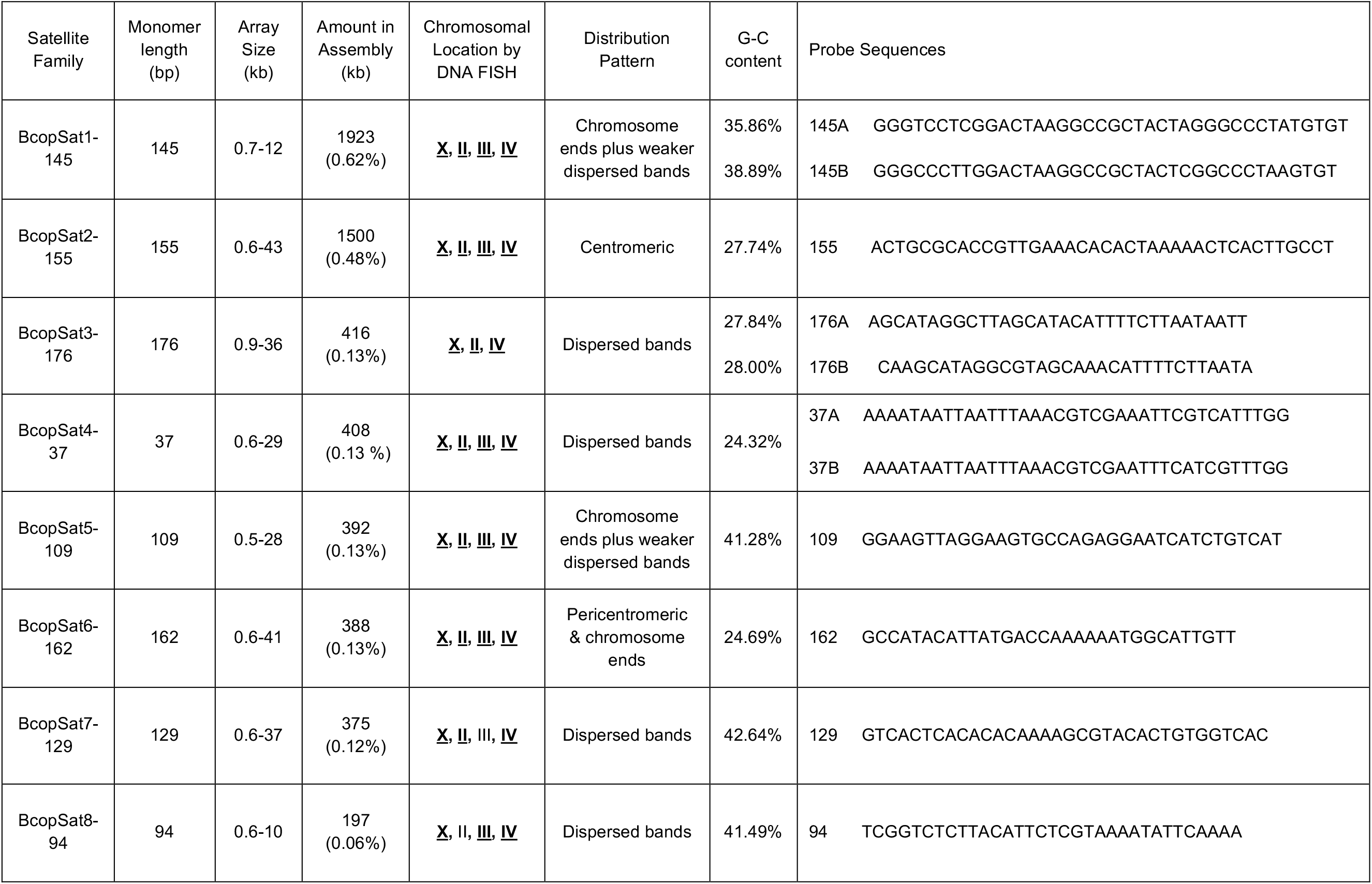
Abundant Satellite Families in *Bradysia coprophila*.

### Cytological mapping of Satellite DNA on the *B. coprophila* polytene chromosomes

To characterize the chromosomal distribution of the newly-identified satellite DNA families, we conducted DNA fluorescence in situ hybridization (FISH) on polytene chromosomes from the salivary glands. Dipteran polytene chromosomes are amplified over 1000 times by endoreplication, allowing the detection of even low abundance DNA sequences that may be undetectable in diploid cells (Zhimulev, 2009). Oligo probes for DNA FISH were designed using conserved DNA sequences within the family, and were thus expected to detect most (if not all) of the satellite DNA variants within each family. For satellite DNAs that had two major variants, namely BcopSat1-145, BcopSat3-176 and BcopSat4-37, we designed probes that are expected to be specific to each variant (Table 1). Indeed, these variant-specific probes were able to discriminate between the two major variants when both probes were used together, although each probe hybridized to both variants when used alone (Figure S3).

DNA FISH to polytene chromosomes confirmed the presence of these satellite DNAs and revealed their distribution along the chromosomes (Table 1, Figures 2-4). We confirmed that BcopSat2-155/Sccr is indeed invariably localized to the centromeres of core chromosomes (Escribá et al., 2011) (Figure 2). We found that a second satellite family, BcopSat6-162, colocalizes with BcopSat2-155 at the centromeres (Figure 2). BcopSat6-162 exhibited additional FISH signals on the distal end of II, and in lower abundance on the other chromosome ends (Figure 2A”, arrowheads).

**Figure 2.**
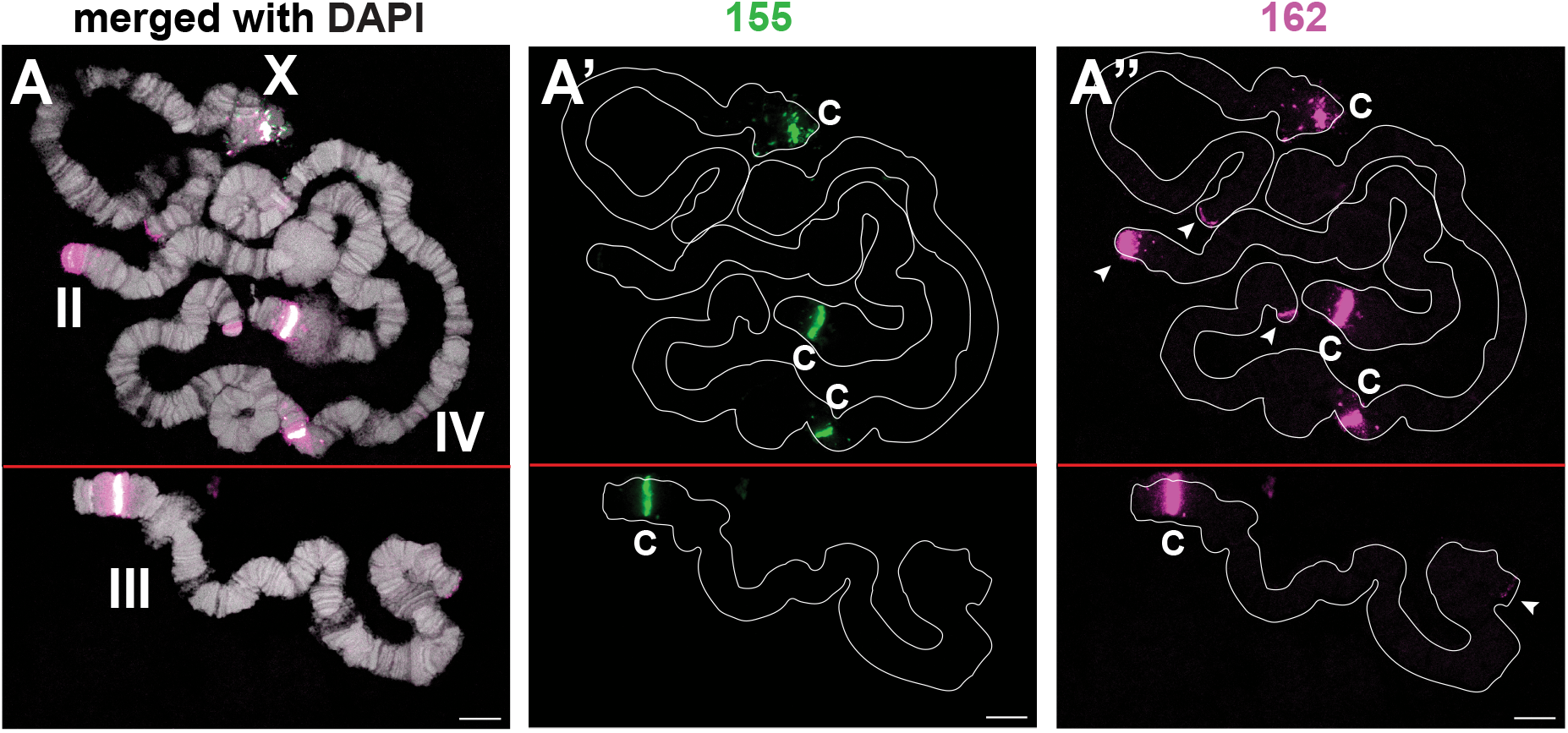
Centromeric and pericentromeric satellite DNA distributions. Distribution of BcopSat2-155 (green) and BcoSat6-162 (magenta) families are shown (A’ and A’’ without DAPI). Both probes hybridize to centromeric and pericentromeric regions, respectively. BcoSat6-162 also hybridizes to telomeric regions of all four chromosomes (arrowheads). Putative centromeres are indicated by “c”. Scale bar represents 10µm.

Other satellite DNAs were distributed outside the centromeres. BcopSat1-145, the most abundant satellite DNA family in the genome, was observed to be dispersed in many shorter arrays along the chromosome arms (Figure 3A). The two variants of BcopSat1-145 (BcopSat1-145A and BcopSat1-145B) were detected at both overlapping and distinct locations along the chromosomes.

**Figure 3.**
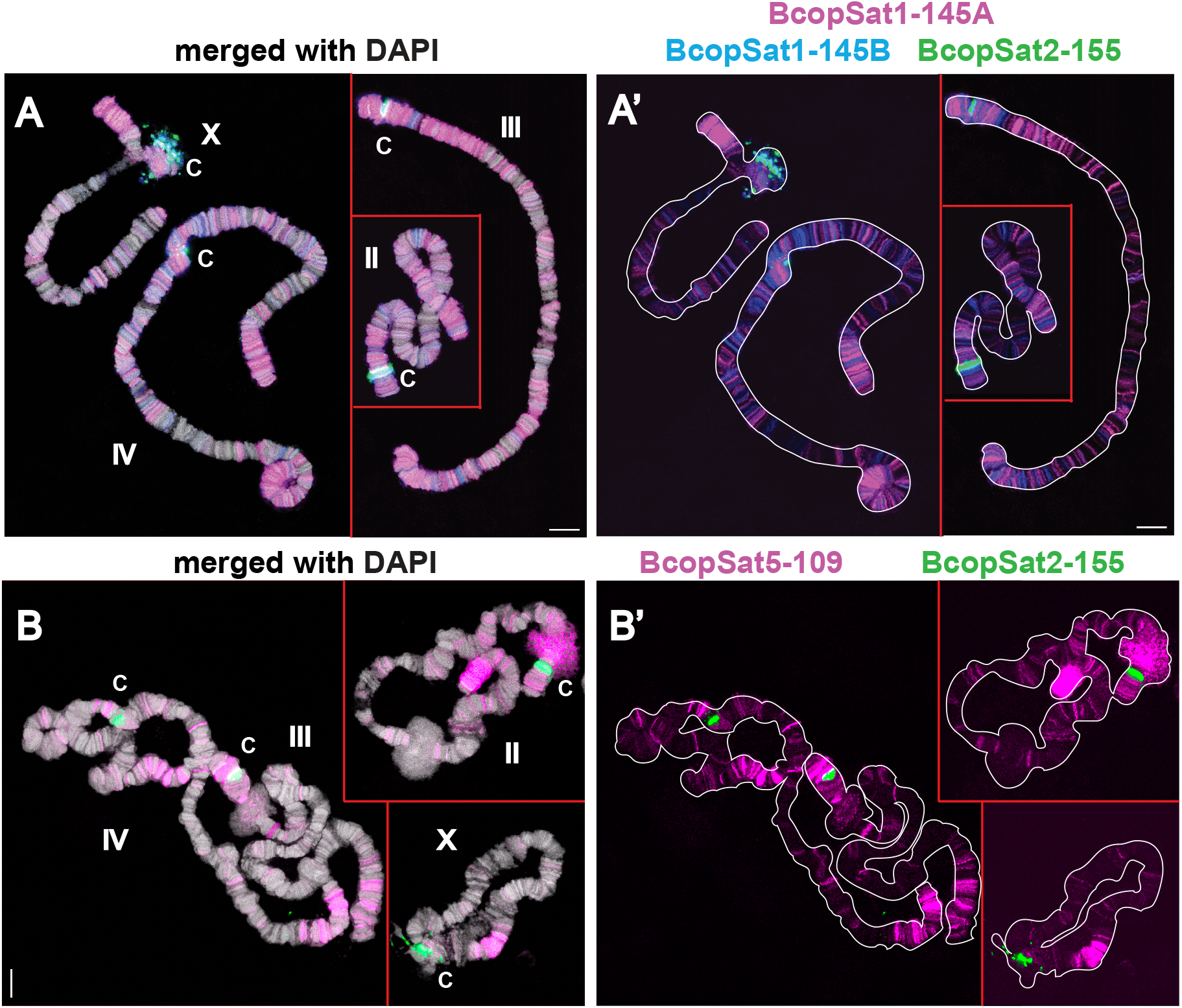
Ubiquitous distributions of satellite DNAs on polytene chromosomes. (A) Hybridization of BcopSat1-145. The 145A (magenta) and 145B (blue) variant sequences are most abundant at the chromosome ends and at the pericentromeric region of chromosome IV (A’ without DAPI). Numerous light bands are present along the arms of all four chromosomes. (B) Hybridization of BcopSat5-109, which is strongest at or near the chromosome ends, with many lighter bands along the chromosome arms (B’ without DAPI). Centromeric regions are shown by BcopSat2-155 (green, c). Scale bar represents 10µm.

BcopSat5-109 was mostly concentrated at or near the chromosome ends with many additional weaker bands along the length of the chromosomes (Figure 3B). BcopSat3-176, BcopSat4-37, BcopSat7-129 and BcopSat8-94 hybridized to fewer discrete bands, often along the chromosome arms (Figure 4). BcopSat3-176A and BcopSat3-176B variant satellite DNAs were found in some non-overlapping bands, suggesting that each variant occupies distinct locations within the genome (Figure 4A). The two variants of BcopSat4-37 (BcopSat4-37A and BcopSat4-37B) rarely overlapped (Figure 4B). BcopSat7-129 families had scattered bands along the arms of the four chromosomes (Figure 4C). The BcopSat8-94 was abundant in blocks in the pericentromeric regions of chromosomes III and IV and in many scattered bands along the chromosome arms (Figure 4D).

**Figure 4.**
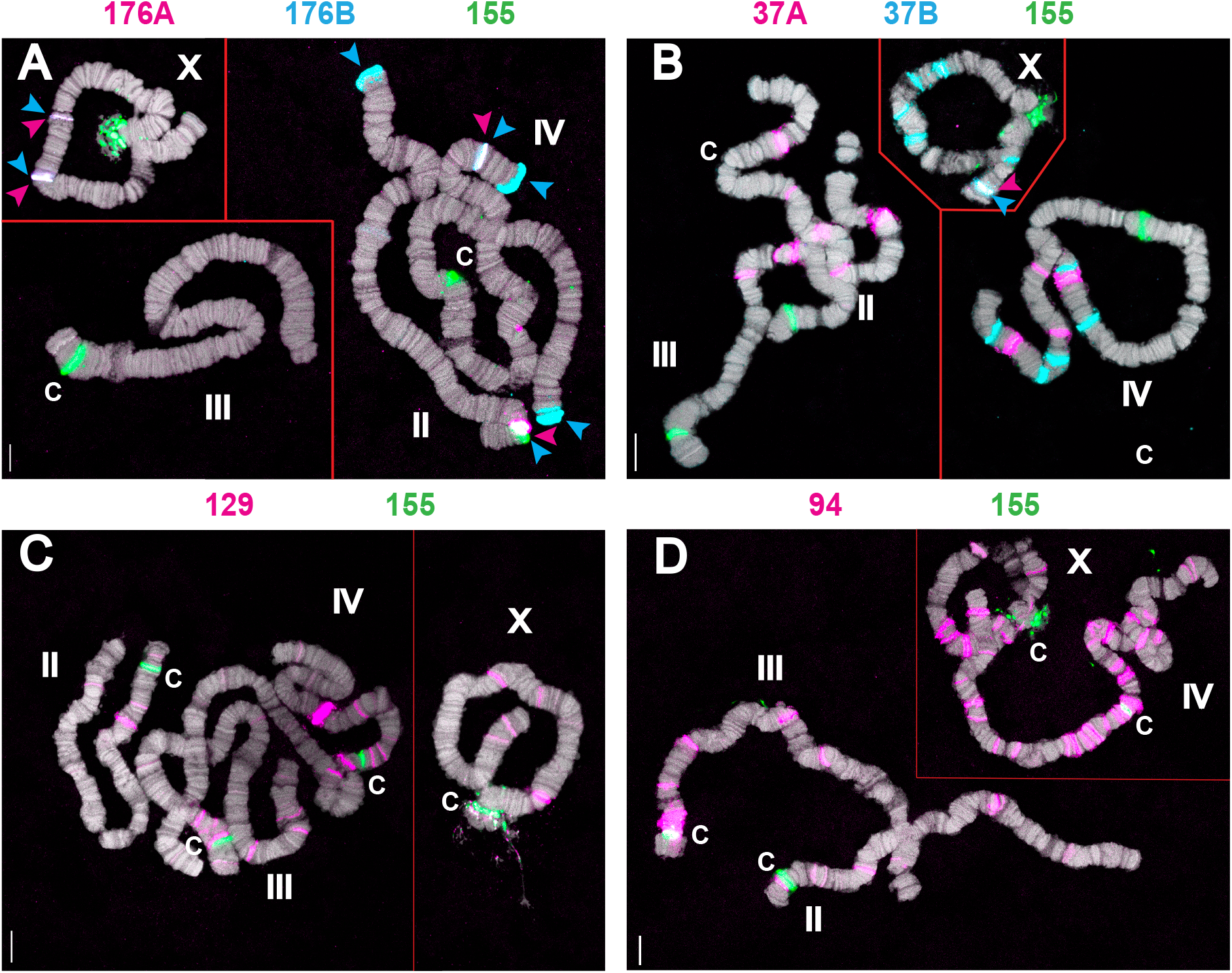
Scattered band distribution of satellite DNA. (A) Hybridization the BcopSat4-176 family. The 176A (magenta) and 176B (blue) probes overlap at all internal bands, but only 176B is found at the chromosome ends. (B) Hybridization of the BcopSat4-37 family. The 37A (magenta) and 37B (blue) probes show that these two variants do not overlap, except for one band on the distal end of the X (blue and magenta arrowheads). (C) Hybridization of the Bcop7-129 family. The 129 probe (magenta) is present in light bands on the arms of all four chromosomes, with one strong band on chromosome IV. (D) Hybridization of the BcopSat8-94 family. The 94 probe (magenta) is abundant in the pericentromeric regions of chromosomes III and IV, and in additional lighter bands on the arms of all four chromosomes. Centromeric regions are shown by BcopSat2-155 (green, c). Scale bar represents 10µm.

### Core chromosomes and L chromosomes have distinct centromeric satellite DNA

Because salivary gland nuclei do not have the germline-limited L chromosomes, we next examined whether the satellite DNA families found on the core chromosomes are also present on L chromosomes using DNA FISH on early pupal testes. The early pupal testis contains cells undergoing meiosis, which can be subjected to chromosome squashes and DNA FISH. L chromosomes in prophase I spermatocytes can be differentiated from X and autosomes based on their ovoid shape (Metz, 1938; Rieffel and Crouse, 1966) (Figure 5A, C). During prophase II, L chromosomes appear as large metacentric chromosomes (Figure 5B, D, E, F).

Although the BcopSat2-155 satellite DNA was previously suggested to present at the centromeres of all chromosomes, including L chromosomes (Escribá et al., 2011), we found that this satellite DNA was present only at the centromeres of the X and autosomes but not on the L chromosomes (Figure 5A, B). The probe sequence used by (Escribá et al., 2011) was cloned using DNA microdissection, and this sequence contained a 78 bp sequence that matches the BcopSat2-155 sequence, as well as a 65 bp sequence from the *B. coprophila* genome that does not match the BcopSat2-155 sequence. This 65 bp sequence appears to be unique to the DNA cloned by (Escribá et al., 2011), but was not associated with any of BcopSat2-155 sequence identified in the genome assembly, perhaps indicating this is a result of cloning artifact (two DNA fragments joined during cloning) or that this chimeric sequence exists in the *B. coprophila* at a very low frequency. To test the possibility that this 65 bp sequence (not present in BcopSat2-155 sequence) might have yielded the FISH signal on L chromosome centromeres in the earlier study (Escribá et al., 2011), we designed multiple probes that cover the entirety of the DNA sequence in the clone described by (Escribá et al., 2011): the 65 bp sequence that does not match BcopSat2-155 sequence, and the 78 bp sequence that matches BcopSat2-155 sequence (Figure S4A). We found that none of probes from the 65bp sequence hybridized to the centromeres of autosomes (Figure S4B,B’, polytene chromosome DNA FISH) or L chromosomes (Figure S4C, meiotic cells DNA FISH), although the signal was detected as multiple faint bands throughout the chromosomes on the polytene chromosomes (Figure S4B,B’). In contrast, the probes against the 78 bp region matching the BcopSat2-155 sequence (155.2, 155.3, Figure S4A) hybridized to the centromeres of core chromosomes, overlapping with the original 155 probe (Figure S4D).

**Figure 5.**
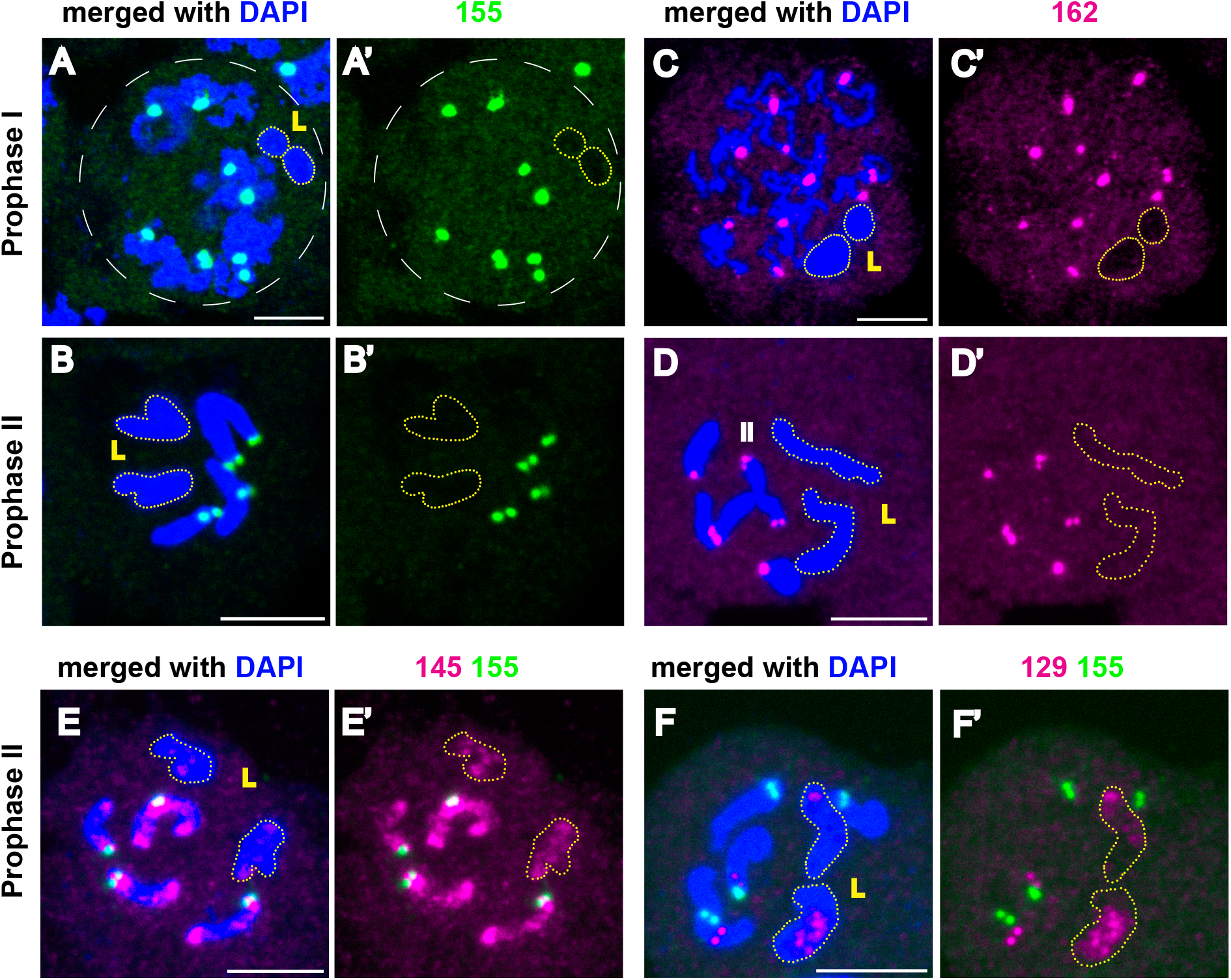
Distributions of satellite DNAs on male meiotic chromosomes. Germline-restricted (L) chromosomes are outlined. Prophase I (A) and prophase II (B) cells with hybridization from BcopSat2-155 (green); prophase I (C) and prophase II (D) cells with hybridization from BcopSat6-162 (magenta). (E) Prophase II cells hybridized with BcopSat2-155 (green) and BcopSat1-145B (magenta). (F) Prophase II cells hybridized with BcopSat2-155 (green) and BcopSat7-129 (magenta). A’-F’ show hybridization in the absence of DAPI. Scale bar represents 5µm.

These data strongly suggest that BcopSat2-155 is only present on the centromeres of the core chromosomes (X and autosomes) and is absent from the L chromosomes. Although the source of the discrepancy between the present work and the previous work by Escriba (Escribá et al., 2011) remains unclear, we note that the previous conclusion that BcopSat2-155 is present on the centromeres of all chromosomes was drawn from the number of foci on chromosome spreads that did not resolve individual chromosomes (Escribá et al. 2011). Thus it is possible that precociously separated sister centromeres or occasional aneuploid cells contributed to their conclusion that BcopSat2-155 is present on the centromeres of core and L chromosomes. Moreover, BcopSat6-162, which we found on the centromeres of X and autosomes (Figure 2), was also not present on the L chromosomes (Figure 5D). These results show that the X chromosome and autosomes do not share centromeric sequence with L chromosomes.

We used probes of the remaining satellite DNA families to the L chromosomes for DNA FISH on meiotic spreads, and found that BcopSat1-145B (Figure 5E), BcopSat1-129 (Figure 5F), and possibly also BcopSat5-109 (Figure S5C), are present on the L chromosomes. Other satellite DNAs that are abundant on the X and autosomes, BcopSat3-176B, BcopSat4-37B, and BcopSat8-94, were not detected on the L chromosomes (Figure S5A, B, D). The 382 bp repeat (BcopSat-382), which was described by (Escribá et al., 2011), showed the expected hybridization patterns (chromosome IV and the L chromosomes) on polytene chromosomes and meiotic spreads (Figure S6).

### Identification of L chromosome-enriched satellite DNA

Considering that the L chromosomes are large heterochromatic chromosomes (Rieffel and Crouse, 1966), it was surprising that most of the abundant satellite DNA families we identified were not detected on the L chromosomes by DNA FISH. Thus, we hypothesized that these chromosomes may contain distinct sets of satellite DNAs compared to the X and autosomes. As the genome assembly that we used to identify *B. coprophila* satellite DNA (Table 1) used male embryos and pupae from stages where most cells have already lost the L chromosomes (Urban et al., 2021), we speculated that L-specific satellite DNA is likely missing from this assembly. We therefore utilized the recent L chromosome assembly, which was generated by comparing somatic (heads) vs. germ (testes) whole genome sequencing (Hodson et al., 2022). Since this assembly was generated using only short read data, we identified 10-fold fewer tandem repeats compared to the core chromosome scaffolds, which were generated from long-read data (Urban et al., 2021). Nonetheless, we were able to identify three abundant satellite DNA repeats with the unit sizes of 10 bp, 38 bp, and 39 bp in the L chromosome assembly (Hodson et al., 2022), which were at very low abundance or absent in the core chromosome scaffolds from the Urban assembly (Urban et al., 2021): BcopSat-10, BcopSat-38 and BcopSat-39 (Table 2).

**Table 2.**
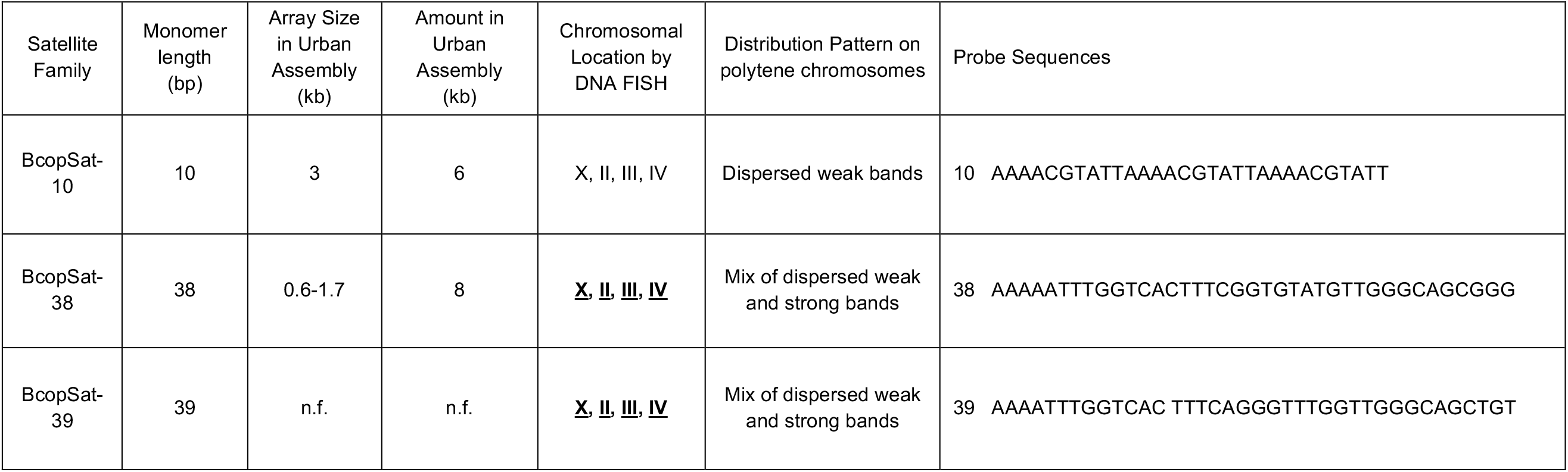
pL satellite families. n.f. = not found.

We conducted DNA FISH on meiotic chromosome spreads to test whether these newly identified satellite DNAs were present on L chromosomes. We found that indeed these three satellite DNAs were enriched on L chromosomes (Figure 6A-C), revealing that unique subsets of satellite DNAs are present on the core vs. L chromosomes. However, these probes (BcopSat-10, BcopSat-38 and BcopSat-39) detected signals at several sites on the X and autosomes of salivary gland polytene chromosomes (Figure S7), suggesting that these satellite DNAs are present on the X and autosomes at low abundance, which becomes only detectable with polytenized chromosomes.

**Figure 6.**
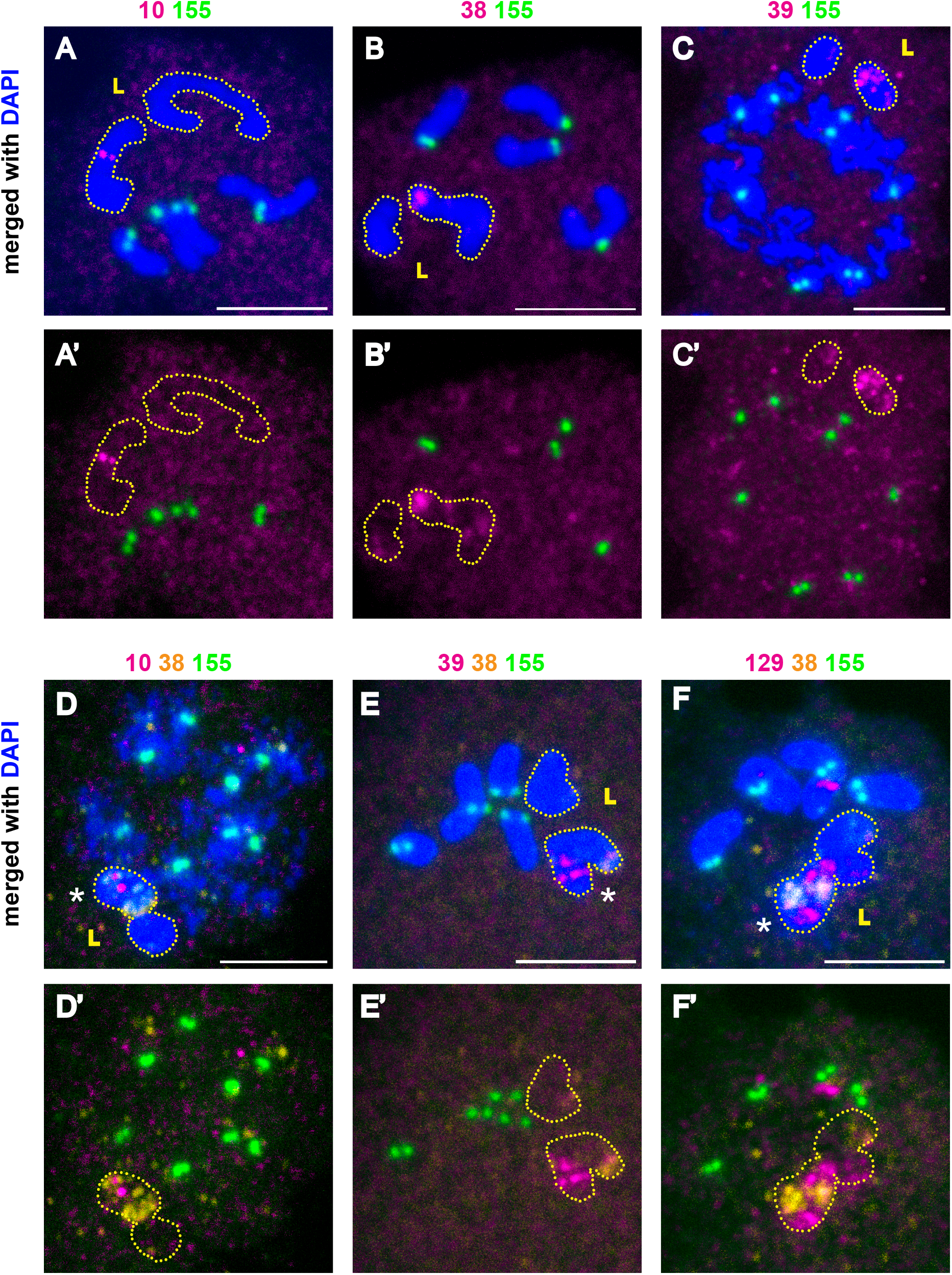
Distributions of other satellite DNAs on male meiotic chromosomes. Germline-restricted (L) chromosomes are outlined. Centromeres are shown by hybridization of BcopSat2-155 (green). (A) Prophase II cells with hybridization from BcopSat-10 (magenta). (B) Prophase II cells with hybridization from BcopSat-38 (magenta). (C) Prophase I cells with hybridization from BcopSat-39 (magenta). (D) Prophase I cells with hybridization from BcopSat-10 (magenta) and BcopSat-38 (yellow). (E) Prophase II cells with hybridization from BcopSat-39 (magenta) and BcopSat-38 (yellow). (F) Prophase II cells with hybridization from BcopSat-129 (magenta) and BcopSat-38 (yellow). Asterisks indicate hybridization to L chromosomes. A’-F’ show hybridization in the absence of DAPI. Scale bar represents 5µm.

### Two L chromosomes in spermatocytes are not homologs

While conducting DNA FISH on L-enriched satellite DNAs (BcopSat-10, BcopSat-38 and BcopSat-39) (Figure 6) as well as BcopSat7-129 (Figure 5F, F’), we were surprised to find that the abundance of these satellite DNAs was not equal between two L chromosomes within a cell. By combining multiple probes (BcopSat-10, BcopSat-38, BcopSat-39 and BcopSat7-129), we found that all of these satellite DNAs were either limited to or enriched on one of the two L chromosomes (Figure 6D-F). This result suggests that there are two distinct L chromosomes, rather than two L homologs, in *B. coprophila* male germline cells. This was surprising because previous cytological studies have assumed that the two L chromosomes found in germ cells were homologs (Metz, 1938). However, a recent study (Hodson et al. 2022) found that the assembled L contigs were larger than the predicted size of a single L chromosome, and there was also bimodality in the coverage of L contigs, implying that the two Ls are distinct chromosomes. Our DNA FISH experiments support this conclusion.

## Discussion

Here we utilized recent genomic data to identify previously unknown satellite DNAs in *B. coprophila*, providing a foundation for future studies on the functions of satellite DNA in this species. We have made two interesting observations regarding the germline-restricted L chromosomes in this species. First, we found that the L chromosomes do not share the centromeric sequences with the core chromosomes (X and autosomes). A similar result has recently been found in two nightingale species, where a candidate centromeric satellite DNA was absent from the germline-restricted chromosome (GRC) (Ridl et al., 2025). It is tempting to speculate that the difference in centromeric satellite DNA may be utilized to allow for L chromosome elimination in somatic cells. For example, sequence-specific DNA binding proteins (specific for X/autosome centromeres, or L centromeres) may promote or interfere with centromere assembly in somatic cells, leading to defective kinetochore function and thus L chromosome elimination. However, cytological studies on *B. coprophila* L chromosome elimination in the soma showed that the L chromosomes appear to correctly separate at the centromeres, but the chromosome ends are not properly separated and the chromosomes are unable to segregate to the poles (de Saint Phalle and Sullivan, 1996; Du Bois, 1933). Such observations are apparently inconsistent with the model that loss of centromere function on the L chromosomes leads to their elimination. It is possible that chromosome elimination during embryonic divisions may involve sequences along the chromosome arms, rather than at the centromeric region. A study looking at the distribution of phosphorylated histone H3S10 during L and X chromosome elimination in early embryos found that there was strong H3S10-P signal on the arms of the chromosomes being eliminated, although H3S10-P was absent from the centromeric regions of the L and X chromosomes and the core chromosomes during elimination (Escribá and Goday, 2013). These results suggests that there are differences in chromatin constitution, which may be involved in the process of chromosome elimination.

In addition to centromeric sequences, our results revealed distinct composition of satellite DNA between the core chromosomes and the L chromosomes. This finding is consistent with the conclusion by Hodson et al. (2022), based on genomic evidence, that the L chromosomes did not arise from polyploidization of the X chromosomes as has been suggested previously (Haig 1993). Based on homologies found between the genes assigned to L chromosome contigs and genes from the species *Mayetiola destructor*, (Hodson et al., 2022) proposed that the L chromosomes entered sciarids via interspecific hybridization with an ancestral member of the Cecidomyiidae family. Identification of satellite DNAs in cecidomyiids in future studies will allow to test this possibility.

Our cytological study supports the conclusion by (Hodson et al., 2022) that there are two distinct L chromosomes, which they called GRC1 and GRC2. Our cytological analysis revealed that multiple satellite DNA probes were more abundant on one L chromosome than the other, demonstrating that satellite DNA compositions are also distinct between two L chromosomes. The finding that there are two kinds of L chromosomes has important implications for the genetics of *Bradysia*. The egg contributes one L chromosome and the sperm two L chromosomes to the zygote, thus the embryo begins with 3 L chromosomes (Figure 1). During early embryogenesis, all three chromosomes are eliminated from somatic cells, but the germ cells also eliminate one of three L chromosomes to have two L chromosomes per germ cell. Earlier studies assumed that two L chromosomes are homologous chromosomes; accordingly, the question ‘which L chromosomes are eliminated in the early germ cells?’ has never been posed. The finding that there are two distinct L chromosomes has new implications for L chromosome elimination. As this study showed, sperm will carry both L1/GRC1 and L2/GRC2 chromosomes. Eggs carry only one L (L1 or L2). Then, the zygote must be L1L1L2 or L1L2L2, and the result of eliminating one L chromosome has three possible outcomes---L1L2, L1L1 or L2L2. In *B. coprophila* and many other sciarid species, females are either exclusively female-producing or male-producing, a phenomenon known as ‘monogenic reproduction’ (Baird et al., 2023). The two female morphs (female-producing vs. male-producing) differ in the presence of a large X-linked inversion (XX’ in female-producing female, XX in male-producing female) (Crouse, 1960). However, it is not currently known whether they also differ in the composition of their L chromosomes. Curiously, male-producing females and males have identical X chromosome composition in their germ cells (both are XX), hinting at the possibility that XX male germ cells and XX female germ cells may have a genetic mechanism to support spermatogenesis vs. oogenesis. It is therefore tempting to speculate that L chromosome composition may be involved in sex determination in the germline, but further work is required to explore this possibility.

In conclusion, we have shown that the core chromosomes and germline-restricted L chromosomes differ in their satellite DNA composition. Notably, the L-chromosomes lack the centromeric satellite DNA sequence found on all core chromosomes, and differ from the core chromosomes in the distribution and abundance of satellite DNA families. One question raised by this study is what is the nature of the centromere on L chromosomes. There are likely L-chromosome specific satellites remaining to be discovered, and possibly one of these defines the L chromosome centromere. We were also able to distinguish two distinct L chromosomes based on their satellite DNA composition. We do not know whether there is functional significance to the presence of two different Ls; however, it is interesting that in males, the L chromosome is the only paternal chromosome that escapes elimination during meiosis I. One possibility is that the two different L chromosomes may be involved in germline sex determination, a question that would be addressed by determining the L chromosome composition in the female germline.

## Materials and Methods

### Fly culture maintenance

The *B. coprophila* HoLo2 and 6980 stocks were obtained from Dr. Susan Gerbi at Brown University. Flies were maintained as described in (Gerbi, 2024).

#### Satellite DNA identification

We used two methods to identify satellite sequences. First, we used Tandem Repeats Finder (TRF,(Benson, 1999)), to identify satellites in the assembly of the core chromosomes (Urban et al. 2021) and the genome assembly that contained L chromosomes (Hodson et al. 2022) using parameters optimized for satellite repeat units of 20-500bp (Figure S1). Second, we searched for simple short (≤ 20 bp) satellites in raw Illumina WGS reads using k-seek (Wei et al., 2014), but using this method we did not identify any such satellites enriched in germline short reads, and could not identify sequences with over 10 kb total abundance in somatic reads. We filtered repeat calls to only those in tandem arrays of at least 5 copies and we standardized the rotation/reverse complement of the consensus sequence of each array. We then used cdhit (Li and Godzik, 2006) to merge related sequences into 191 clusters representing different satellite families. We selected the eight most abundant satellite families (present at ≥200 kb in the assembly) for further analysis. We generated a BED file of all contigs containing tandem repeats from the raw TRF output to analyze the locations and abundances of the 8 repeats selected.

### DNA FISH

DNA FISH was performed to chromosome squashed using a modified protocol of (Larracuente and Ferree, 2015). Sample preparation: For salivary gland polytene chromosome preparation, salivary glands were dissected from 4^th^ instar larvae in 45% acetic acid and moved to 12 μl of 45% acetic acid on a clean Superfrost slide (VWR). A coverslip was added and the sample was gently tapped to spread the chromosomes. A rubber stamp was then used to manually squash the preparation, and the sample was frozen by submerging it in liquid nitrogen. Subsequently, the coverslip was removed and the slides were washed in 100% ethanol and air-dried. Slides were fixed in 3.7% formaldehyde in PBS for 4 minutes at room temperature. The slides were washed in PBS, 2XSSC, and dehydrated in 70% ethanol and 100% ethanol. Slides were store in a dark, dust-free location until hybridization. For meiotic chromosome spreads, testes were dissected from young pupae (1/3-1/2 filled eyespots) in PBS. The testes were moved to 20 μl of 2.2% formaldehyde in 45% acetic acid, and fixed for 4 minutes at room temperature. A coverslip was placed onto the sample, and the sample was squashed by applying the rubber stamp onto it. The sample was then frozen in liquid nitrogen, and the coverslip was removed. The slide was rinsed in 100% ethanol, and air-dried. Prior to hybridization, the sample was treated with RNAaseA (2mg/ml) for 10 minutes at 37°C, and subsequently washed in PBS, and dehydrated in 70% ethanol then 100% ethanol, and air-dried. Slides were stored as above. Hybridization: Oligos (30-40 mer) labeled with Alexa-488, Cy3 or Cy5 were used as probes (probe sequences are provided in Table 1). 20 μl of hybridization buffer (50% formamide, 10% dextran sulfate, 2X SSC and 0.5 mM of each probe) was added onto the dry sample, and a coverslip was placed onto the sample. The sample was treated either at 91°C for 3 minutes (salivary glands) or at 95°C for 5 minutes (testes squashes) to denature DNA. The slides were cooled briefly, covered with parafilm and placed in a humid chamber in the dark, and incubated overnight at room temperature. The next day, the coverslips were removed, and the slides were washed at room temperature in 2X SSC (two times for 10 minutes) followed by 0.2X SSC (once for 10 minutes) for salivary glands, or in 0.1X SSC (three times for 15 minutes) for testes squashes. The slides were air-dried in the dark, and mounted in the Vectashield mounting medium containing DAPI. Images were taken using a SP8 confocal microscope with a 63X oil immersion objective and processed using ImageJ software (Fiji).

