## Supplementary information for "Distinct satellite DNA composition between core and germline restricted chromosomes in *Bradysia (Sciara) coprophila*"

### Supplementary figures

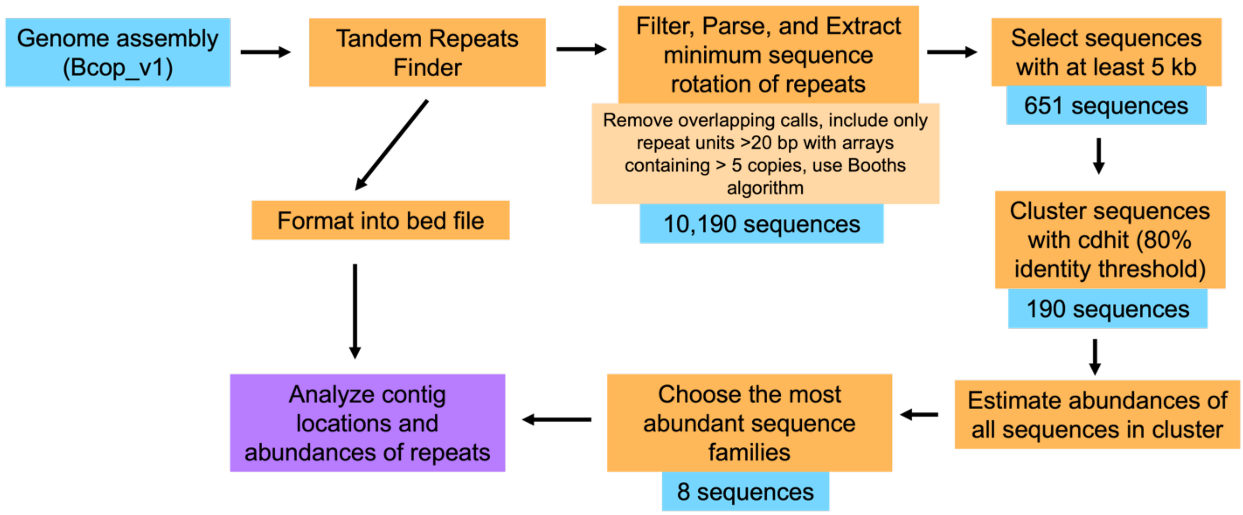


**Figure S1. Satellite repeat identification flowchart.**

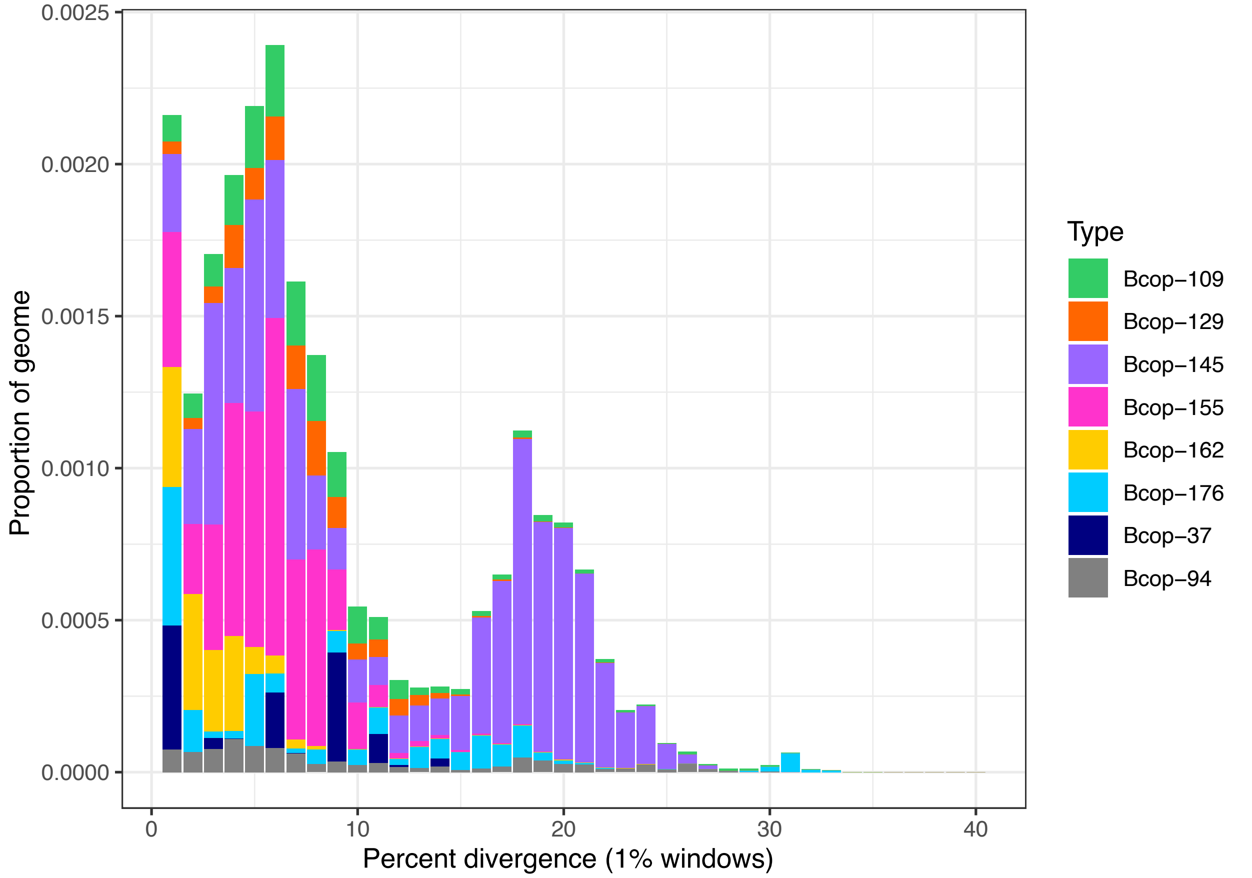


**Figure S2. Repeat divergence landscape.** Each satellite DNA family is indicated by a different color. Divergence from a consensus sequence is shown along the x-axis, and abundance of sequence variants is shown along the y-axis. The data for this plot was produced by running RepeatMasker against the genome assembly using consensus sequences of the 8 satellite families as the library, followed by using the following script: https://github.com/4ureliek/Parsing-RepeatMasker-Outputs/blob/master/parseRM.pl. For satellite families BcopSat4-37and BcopSat1-145, we can see multiple discrete peaks representing abundant sequence variants within the family.

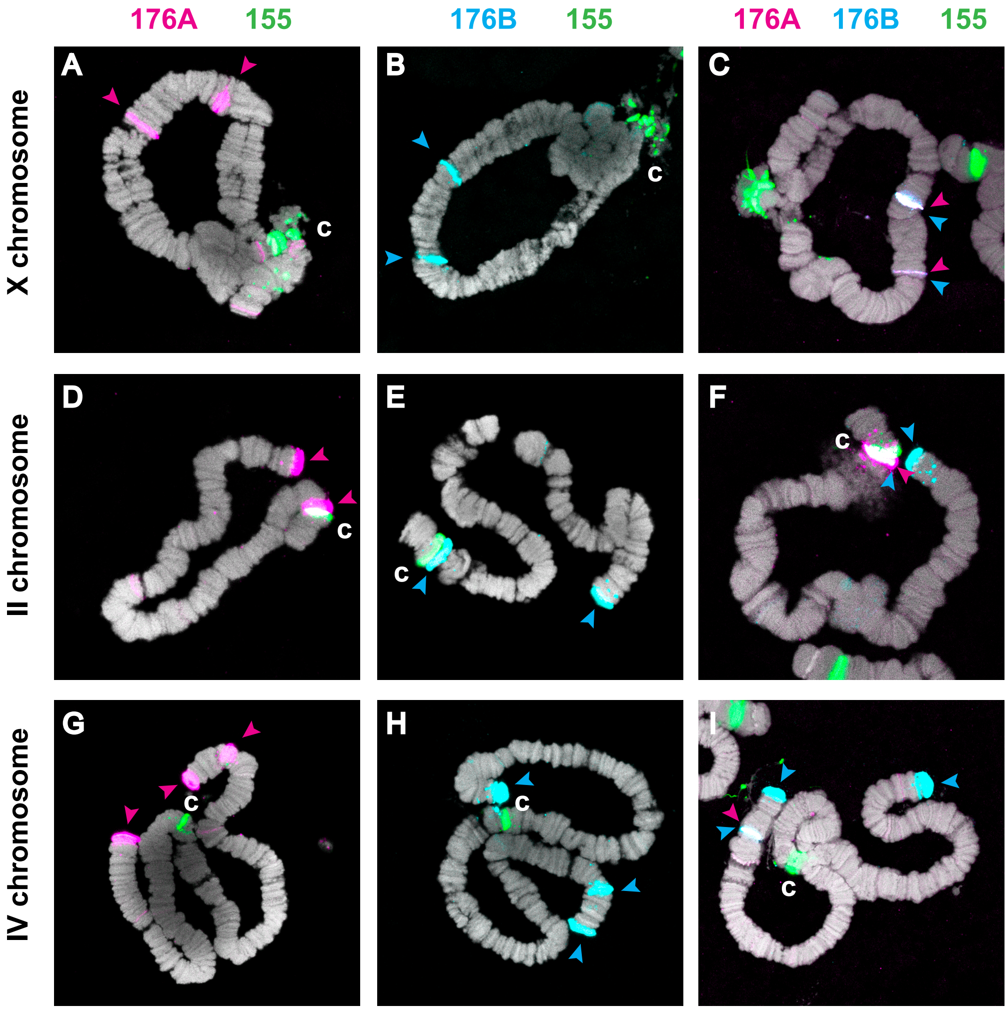


**Figure S3. Cross-hybridization of 176A and 176B probes.** Centromeric regions (c) in all panels are shown by hybridization of BcopSat2-155 probe (green) (A,B,C) Hybridization to the X chromosome. (A) The BcopSat3-176A probe used alone and (B) the BcopSat3-176B probe used alone recognize two bands on the X chromosome which overlap when both probes are used (C). (D,E,F) Hybridization to chromosome II. (D)The BcopSat3-176A probe used alone and the (E) BcopSat3-176B probe used alone hybridize to a band near the centromere and at the distal end when used alone, but only the BcopSat3-176B probe is found at the distal end when both probes are used (F). (G,H,I) Hybridization to chromosome IV. (G)The BcopSat3-176A probe used alone and the (H) BcopSat3-176B probe used alone hybridize to a three bands at or near the ends of the chromosome, whereas only the BcopSat3-176B probe hybridizes to the bands at the chromosome ends when both probes are used (I).

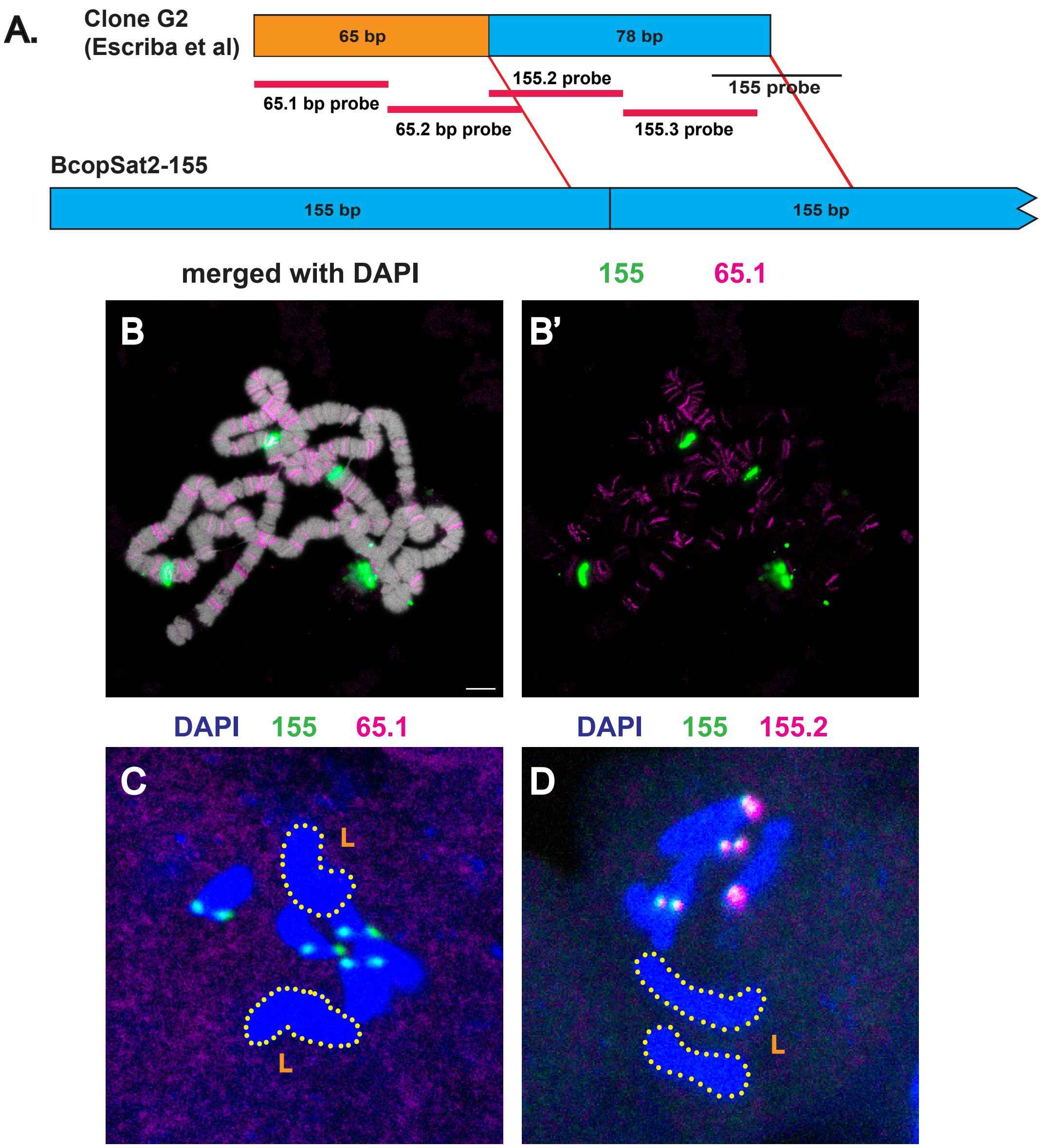


**Figure S4.** (A) Arrangement of sequences within the insert, showing the placement of non-155 sequences (orange, 65bp adjacent to 78bp of sequences (blue) from the 155/Sccr repeat). (B) Hybridization of 155bp sequences (155bp probe, green) and 65bp sequences (65bp probe, magenta) to polytene chromosomes. (C) Hybridization of 155bp (green) and 65bp (magenta) sequences (probe 65.1) to a prophase II cell. (D) Hybridization showing colocalization of the original 155 probe (green) and one of the new 155 probes (155.2) to a prophase II cell.

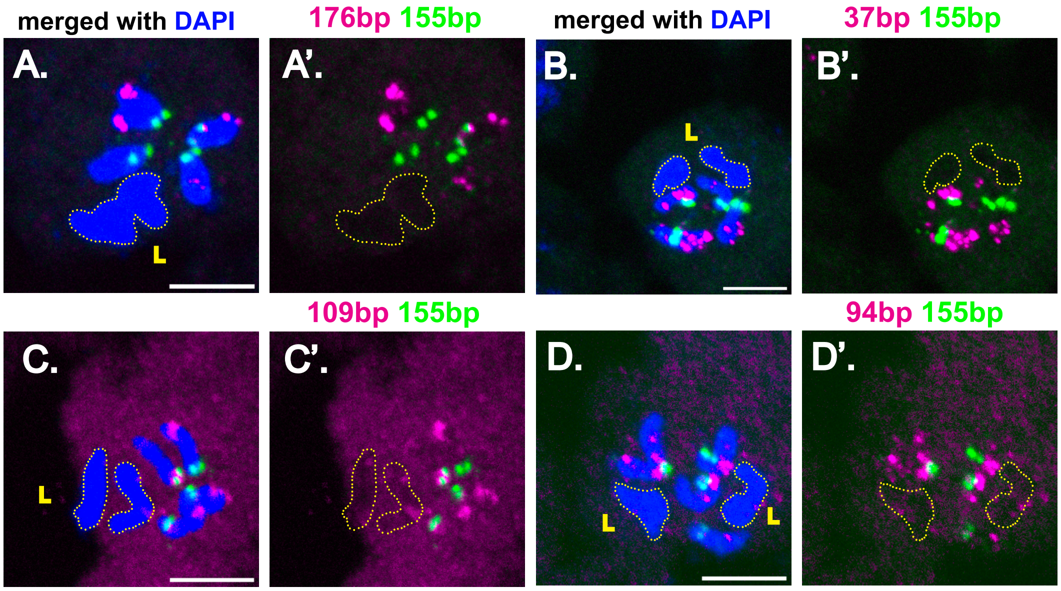


**Figure S5. Hybridization of satellite families to prophase II meiotic chromosomes.** Hybridization with probes to BcopSat3-176B (A, magenta), BcopSat4-37B (B, magenta), BcopSat5-109 (C, magenta), and BcopSat8-94 (D, magenta) are shown. X chromosome and autosome centromeres in all panels are shown by hybridization of the BcopSat2-155 probe (green). (A'-D') show hybridization without DAPI. Germline-restricted (L) chromosomes are outlined. Scale bar represents 5μm.

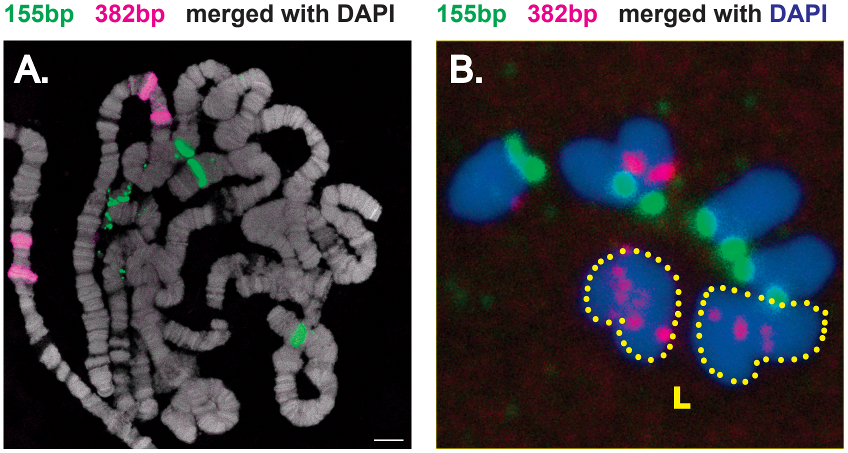


**Figure S6. Hybridization of 382 bp sequences to polytene and meiotic chromosomes.** (A) 382 bp sequences (magenta) are found within two bands on one arm of chromosome IV. We did not detect hybridization of this probe to the proximal heterochromatin of the X chromosome. (B) Prophase II cell. The 382 bp sequences (magenta) also hybridize to both L chromosomes. Hybridization to one arm of the metacentric chromosome IV can also be seen. Centromeres of the X and autosomes are marked by hybridization of the BcopSat2-155 probe (green).

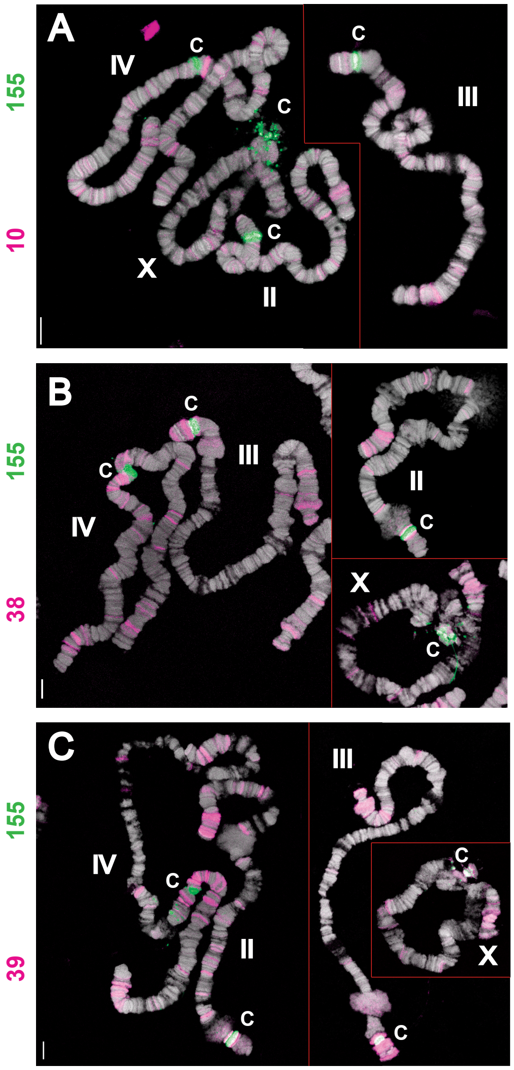


**Figure S7. DNA FISH of L chromosome satellites to polytene chromosomes.** Hybridization of BcopSat2-10 (A, magenta), BcopSat-38 (B, magenta) and BcopSat-39 (C, magenta) shown. A-C are composite figures. Centromeres in all panels are shown by hybridization of the BcopSat2-155 probe (green) and are indicated by "c". Scale bar represents 10µm.
